# Among-individual diet variation within a lake trout ecotype: lack of stability of niche use

**DOI:** 10.1101/811851

**Authors:** L. Chavarie, K.L. Howland, L.N. Harris, C.P. Gallagher, M.J. Hansen, W.M. Tonn, A.M. Muir, C.C. Krueger

## Abstract

In a polymorphic species, stable differences in resource use are expected among ecotypes, and homogeneity in resource use is predicted within an ecotype. Yet, using a broad resource spectrum has been identified as a strategy for fishes living in unproductive northern environments, where food is patchily distributed and ephemeral. We investigated whether individual specialization of trophic resources occurred within the generalist piscivore ecotype of lake trout from Great Bear Lake, Canada, reflective of a form of diversity. Four distinct dietary patterns of resource use within the lake trout ecotype were detected from fatty acid composition, with some variation linked to spatial patterns within Great Bear Lake. Feeding habits of different groups within the ecotype were not associated with detectable morphological or genetic differentiation, suggesting that behavioral plasticity caused the trophic differences. A low level of genetic differentiation was detected between exceptionally large-sized individuals and other individuals. Investigating a geologically young system that displays high levels of intraspecific diversity and focusing on individual variation in diet suggested that individual trophic specialization can occur within an ecotype. The characterization of niche use among individuals, as done in this study, is necessary to understand the role that individual variation can play at the beginning of differentiation processes.

## Introduction

Phenotypic diversity within fish species that have colonized post-glacial lakes often re-present early stages of species diversification (Snorrason et al., 2004). Many fishes that have colonized post-glacial freshwater systems are assumed to have been plastic generalists (i.e., flexible in use of habitat and food resources) at the time of colonization (Skúlason et al., 2019; Snorrason et al., 2004). Given the novel environment and new ecological opportunities, a newly established population may begin to display among-individual differences in behavior and other phenotypic characteristics (Svanbäck et al., 2007). Phenotypic plasticity, the capacity for one genotype to produce different phenotypes in response to environmental cues, could be a character subject to selection, facilitating the process of diversification (De Jong, 2005). Despite uncertainties of how phenotypic plasticity promotes divergence, plasticity appears to serve as an important element in early phases of diversification (Handelsman et al., 2013; Nonaka et al., 2015; Snorrason et al., 2004). Theory predicts that stable and predictable recently colonized systems would favor foraging and habitat specialization and increase the probability of eco-morphological diversification (Skúlason et al., 1999; Snorrason et al., 2004; Van Kleunen et al., 2005).

Phenotypic plasticity in temporally and spatially variable environments has been demonstrated repeatedly within and among populations (Skúlason et al., 2019). Whether niche expansion of a population is achieved by a general increase in niche widths for all individuals overall or by an increase of among-individual variation (i.e., expression of multiple individual specializations within a population) is a question in evolutionary ecology that remains unanswered (Bolnick et al., 2003; Roughgarden, 1972; Svanbäck et al., 2012). Several apparent generalist populations have been reported to be composed of combinations of specialized individuals using several narrow niches that together yield an overall wide population niche (Araújo et al., 2011; Araújo et al., 2008; Bolnick et al., 2003). Post-glacial lakes and coinhabiting species offer a wide range of characteristics that may favor or constrain individual specialization. Post-glacial lakes are depauperate ecosystems with low interspecific competition, which provides ecological opportunities that likely favor niche expansion (Bolnick et al., 2010; Costa et al., 2008; Parent et al., 2014). Additionally, the large flexibility within post-glacial colonizing species, with individuals having the potential to exploit a wide range of resources, can facilitate the evolution of individual resource specialization and population divergence. Yet, northern ecosystem food-webs are subjected to strong seasonal and episodic influences of climate and the environment (McMeans et al., 2015). Accordingly, using a broad resource spectrum has been identified as a useful strategy for fishes living in Arctic environments, where food can be patchily distributed and ephemerally available. From all the facets of niche use that are possible in northern lakes, understanding the magnitude and effect of individual specialization in species and trophic positions is necessary to understand the role that variation among individuals can play at the beginning of differentiation processes (Cloyed et al., 2016; De León et al., 2012; Svanbäck et al., 2015).

Great Bear Lake (Northwest Territories, Canada), straddling the Arctic Circle, provides an excellent opportunity to investigate the role of among-individual diet variation in diversification processes in post-glacial lakes. Lake trout, *Salvelinus namaycush*, in this lake show a high degree of intraspecific diversity within a geologically young system (8,000–10,000 yr BP; Johnson, 1975; Pielou, 2008). Specifically, extensive sympatric divergence has occurred for this species with four ecotypes inhabiting the shallow-water (≤ 30 m) zone of Great Bear Lake (Fig. A1; Chavarie et al., 2016a; Chavarie et al., 2015; Chavarie et al., 2013; Harris et al., 2015). Three of these four shallow-water lake trout ecotypes are described as trophic generalists with differing degrees of omnivory along a weak benthic-pelagic gradient (Chavarie et al., 2016a; Chavarie et al., 2016b). Despite habitat and dietary overlap, significant differences in morphological, genetic, and life-history variation have been reported (Chavarie et al., 2016; Chavarie et al., 2013; Harris et al., 2015). The suggested resource use among the three ecotypes could be caused by the combination of individual specialists along a resource continuum (Chavarie et al., 2016b). In other words, although ecotype resource use may appear similar, individuals within an ecotype may differ in their resource use. One of these three generalist ecotypes (Ecotype 2; generalist with a tendency to consume more fish than other ecotypes, referred to here as the piscivorous ecotype; Fig. 1) showed at least two different feeding strategies, benthic cannibalism and interspecific piscivory in the pelagic zone (Chavarie et al., 2016c).

**Fig. 1.**
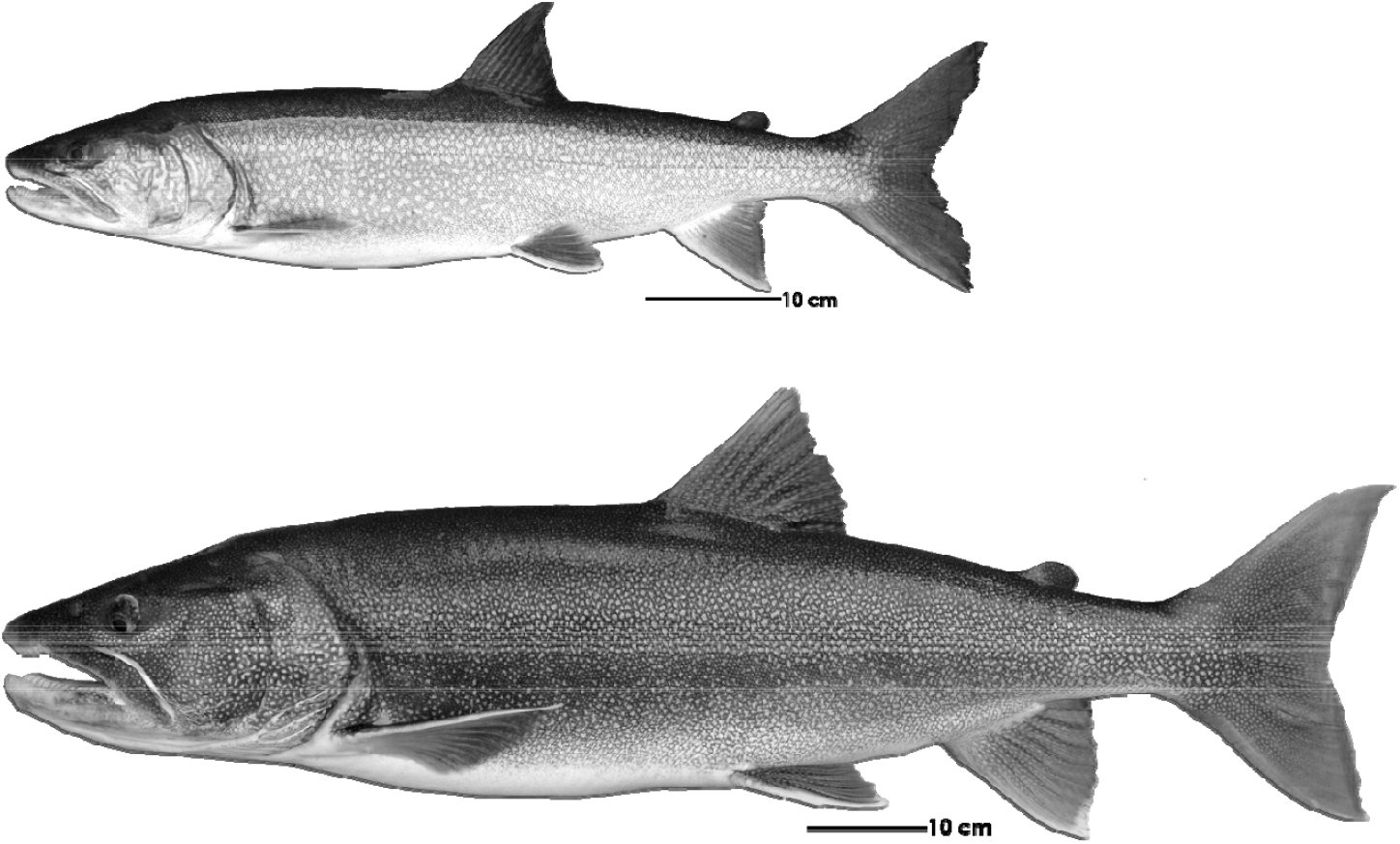
Example of a piscivorous (64 cm) and a Giant (100 cm standard length) Lake Trout, respectively, from Great Bear Lake (NT).

To characterize niche use and individual variation within an ecotype in relation to observed differentiation of feeding strategies, we focused this study solely on the piscivorous lake trout ecotype. Fatty acid analysis assumes that dietary lipids are broken down into their constituent fatty acids and incorporated relatively unchanged into consumer tissues (Howell et al., 2003; Iverson, 2009; Iverson et al., 2004), allowing spatial and temporal diet comparison among individuals (Duerksen et al., 2014; Eloranta et al., 2011; Hoffmann, 2017; Iverson, 2009; Scharnweber et al., 2016). Fatty acids have been assessed to be a robust tool to characterize lake trout diets (Happel et al., 2017; Happel et al., 2016; Iverson, 2009). Thus, fatty acids were used as trophic bio-indicators to better understand dietary patterns of piscivorous lake trout and investigate whether variation occurred among individuals in this ecotype and if individual specialization may be contributing to the trophic breadth of the ecotype. Specifically, our aims were to 1) compare resource use among lake trout individuals within Ecotype 2 (piscivores) by characterizing their fatty acids profiles, 2) determine whether resource-use differences were influenced by life-history traits (e.g., size and age), 3) characterize and compare morphological variation among groups that expressed different feeding strategies, and 4) determine if genetic differences existed among groups. In addition, we examined a sub-set of large lake trout of this ecotype from our collections (> 900 mm in fork length) referred to locally as “Giants” (Fig. 1), to determine if they showed any ecological and genetic differences from others of this ecotype. These exceptionally large individuals comprise < 1% of the lake trout population sampled in Great Bear Lake, and are among the largest lake trout in the world (Chavarie et al., 2016). Except for their large body-size, these individuals show no major morphological or spatial and temporal distribution differences relative to other co-occurring piscivorous lake trout.

## Methods

### Study area and field sampling

Great Bear Lake is an oligotrophic Arctic freshwater system, 250 km south of the Arctic Ocean, in Northwest Territories, Canada (N66° 06’ W120° 35’) (Johnson, 1975). As the world’s ninth largest and 19^th^ deepest lake, the lake has a complex, multi-armed surface area of 31,790 km^2^ and a maximum depth of 446 m (mean depth = 90 m). Great Bear Lake was formed by scouring from the Laurentide ice-sheet during the Pleistocene and was originally part of glacial Lake McConnell 8,000–10,000 yr BP (Johnson, 1975; Pielou, 2008). The lake has characteristics typical of an arctic lake: ultra-oligotrophic, short ice-free season, and a simple food web supporting only 15 fish species (Alfonso, 2004; Johnson, 1975; MacDonald et al., 2004). Great Bear Lake lacks a commercial fishery but plays an important role in the local economy, supporting a fly-in sport fishery for tourists and a subsistence fishery for the small Sahtu community of Déline. Great Bear Lake has considerable intraspecific diversity within lake trout, lake whitefish *(Coregonus clupeaformis)*, and cisco *(C. artedi)* (Chavarie et al., 2013; Howland et al., 2013).

Piscivorous lake trout were caught at depths ≤ 30 m using paired bottom sets (ca. 24 h) of 140-mm and multi-mesh (38–140 mm) stretched-mesh gill nets from late-July through August over multiple years (2002–2011) among all five arms of the lake (Chavarie et al., 2016b; Chavarie et al., 2015; Chavarie et al., 2013). During 2012-2014, multi-mesh gill nets (38 to 140 mm), with a typical soak time of 24 hours, were distributed across random depth-stratified sites (0–150 m) among Keith, McVicar, and McTavish arms (Table A1). Compared to the other ecotypes, piscivores have a streamlined body, large gape, and high growth rates throughout life, similar to other piscivores (Chavarie et al., 2016; Chavarie et al., 2013). The piscivorous ecotype also displayed a modest level of genetic differentiation from the three other ecotypes (Harris et al., 2015).

We focused on adult trout due to the difficulty of classifying juveniles into ecotypes (Chavarie et al., 2013; Zimmerman et al., 2006; Zimmerman et al., 2007) and to avoid the confounding effects of ontogenetic shifts in morphology and diet. Of 79 fish analyzed herein, 35 piscivourous lake trout (Ecotype 2) were previously analyzed for fatty acids by Chavarie et al. (2016b) and 44 fish were new additions to the diet analyses presented here. Fish were selected from collections analyzed morphologically by Chavarie et al. (2015) to include a range of sizes and ages within the piscivorous ecotype. For analyses invloving giant individuals, we selected lake trout with fork lengths > 900 mm.

A left lateral full-body digital image was taken for each lake trout caught according to the procedures in Muir et al. (2012). Measurements, tissues, and structures were sampled to determine biological characteristics related to life-history, including otoliths, fork length, round weight, sex, and stage of maturity (i.e., immature, current year spawner, or resting) (Chavarie et al., 2016; Chavarie et al., 2013). A dorsal muscle sample was collected and frozen at −20°C for fatty acid analysis (Budge et al., 2006; Kavanagh et al., 2010; Loseto et al., 2009) and tissue from pectoral fins was collected and preserved in 95% ethanol for genetic analyses.

### Fatty Acids

Analysis of 41 dietary fatty acids was carried out using procedures described by Chavarie et al. (2016b) (Table 1). Muscle samples were freeze-dried and subsequently homogenized with a mortar and pestle. Lipids were extracted overnight from 1 g of the homogenate in a 2:1 chloroform-methanol solution containing 0.01% BHT (v/v/w) at −20°C (Folch et al., 1957). After extraction, samples were filtered through Whatman Grade 1 Qualitative filter paper and the filter paper/sample was rinsed twice with 2 ml of the 2:1 chloroform:methanol. Sample extract was collected in a test tube and 7 ml of 0.88 N NaCl solution was added to encourage fatty acids to move into the organic (chloroform) layer. The aqueous layer was discarded after which the chloroform was dried with sodium sulfate prior to total lipid measurement. The extracted lipid was used to prepare fatty acid methyl esters (FAME) by transesterification with Hilditch reagent (0.5 N H_2_SO_4_ in methanol) (Morrison et al., 1964). Samples were heated for 1 h at 100 °C. Gas chromatographic (GC) analysis was performed on an Agilent Technologies 7890N GC equipped with a 30 m J&W DB-23 column (0.25 mm I.D; 0.15 μm film thickness). The GC was coupled to a Flame Ionization Detector operating at 350 °C. Hydrogen was used as carrier gas flowing at 1.25 ml/min for 14 minutes, and increased to 2.5 ml/min for 5 min. The split/splitless injector was heated to 260 °C and run in splitless mode. The oven program was as follows: 60 °C for 0.66 min, increasing by 22.82 °C/min to 165 °C with a 1.97 min hold; increasing by 4.56 °C/min to 174 °C and by 7.61 °C/min to 200 °C with a six min hold. Peak areas were quantified using Agilent Technologies ChemStation software. Fatty acids standards were obtained from Supelco (37 component FAME mix) and Nuchek (54 component mix GLC-463).

**Table 1.** Mean composition (% ± SD) of 41 fatty acids for the four groups of piscivorous Lake Trout morph identified from Great Bear Lake.

| **Fatty acids** | **Group 1** | **Group 2** | **Group 3** | **Group 4** |
| --- | --- | --- | --- | --- |
| 14:0 | 6.8 ± 1.0 | 7.0 ± 1.0 | 9.2 ± 1.0 | 9.9 ± 1.1 |
| 16:0 | 28.1 ± 1.0 | 28.13 ± 1.7 | 24.2 ± 1.7 | 26.1 ± 1.2 |
| 16:1n-7 | 15.9 ± 3.7 | 10.1 ± 2.0 | 19.5 ± 3.4 | 15.6 ± 2.1 |
| 16:2n-6 | 2.0 ± 0.5 | 2.4 ± 0.6 | 2.6 ± 0.2 | 3.1 ± 0.2 |
| 16:2n-4 | 2.6 ± 0.7 | 1.5 ± 0.4 | 2.7 ± 0.9 | 2.3 ± 0.4 |
| 17:0 | 2.7 ± 0.5 | 2.8 ± 0.3 | 2.4 ± 0.4 | 2.8 ± 0.2 |
| 16:3n-4 | 1.5 ± 0.7 | 1.4 ± 0.5 | 1.9 ± 0.6 | 1.8 ± 0.9 |
| 16:4n-3 | 2.6 ± 1.2 | 0.8 ± 0.3 | 1.3 ± 0.6 | 1.2 ± 0.4 |
| 16:4n-1 | 1.6 ± 0.7 | 1.5 ± 0.8 | 0.9 ± 0.6 | 1.0 ± 0.6 |
| 18:0 | 14.2 ± 1.6 | 13.1 ± 0.8 | 11.6 ± 0.7 | 11.7 ± 0.5 |
| 18:1n-9 | 20.6 ± 4.1 | 18.5 ± 3.4 | 32.3 ± 3.9 | 27.9 ± 3.5 |
| 18:1n-7 | 11.9 ± 2.4 | 9.5 ± 1.0 | 13.9 ± 1.2 | 12.5 ± 0.8 |
| 18:2n-6 | 8.6 ± 1.5 | 9.2 ± 1.6 | 12.4 ± 1.2 | 12.9 ± 1.0 |
| 18:2n-4 | 2.0 ± 0.4 | 1.5 ± 0.2 | 2.1 ± 0.2 | 2.1 ± 0.2 |
| 18:3n-6 | 2.2 ± 0.8 | 1.5 ± 0.4 | 2.5 ± 0.4 | 2.3 ± 0.2 |
| 18:3n-4 | 2.2 ± 0.7 | 1.5 ± 0.3 | 2.4 ± 0.4 | 2.0 ± 0.3 |
| 18:3n-3 | 6.6 ± 1.4 | 6.9 ± 0.9 | 7.9 ± 0.6 | 8.7 ± 0.7 |
| 18:3n-1 | 1.2 ± 0.7 | 1.2 ± 0.3 | 1.1 ± 0.3 | 1.5 ± 0.3 |
| 18:4n-3 | 3.5 ± 0.7 | 4.0 ± 1.2 | 4.9 ± 0.7 | 5.6 ± 0.7 |
| 18:4n-1 | 1.3 ± 0.6 | 0.4 ± 0.5 | 0.9 ± 0.5 | 1.2 ± 0.6 |
| 20:0 | 2.1 ± 0.7 | 2.8 ± 0.7 | 3.1 ± 0.6 | 2.8 ± 0.8 |
| 20:1n-11 | 1.7 ± 1.0 | 0.8 ± 0.5 | 1.9 ± 0.8 | 1.4 ± 0.4 |
| 20:1n-9 | 6.0 ± 1.4 | 4.2 ± 0.8 | 7.9 ± 0.9 | 7.1 ± 0.9 |
| 20:1n-7 | 2.5 ± 0.4 | 2.5 ± 0.3 | 3.8 ± 0.4 | 4.1 ± 0.6 |
| 20:2n-9 | 0.8 ± 0.6 | 1.4 ± 0.8 | 1.3 ± 0.4 | 1.2 ± 0.4 |
| 20:2n-6 | 3.8 ± 0.9 | 4.7 ± 0.9 | 6.8 ± 1.3 | 7.5 ± 1.0 |
| 20:3n-6 | 3.4 ± 0.5 | 3.6 ± 0.4 | 4.4 ± 0.5 | 4.0 ± 0.4 |
| 20:4n-6 | 13.8 ± 1.7 | 14.2 ± 1.3 | 10.1 ± 1.1 | 10.0 ± 1.2 |
| 20:3n-3 | 3.5 ± 0.7 | 4.5 ± 0.9 | 5.1 ± 0.6 | 6.6 ± 0.7 |
| 20:4n-3 | 6.1 ± 1.2 | 8.2 ± 1.3 | 8.8 ± 1.1 | 10.8 ± 0.9 |
| 20:5n-3 | 18.0 ± 2.9 | 15.7 ± 1.2 | 11.8 ± 2.1 | 12.2 ± 1.8 |
| 22:1n-11 | 1.8 ± 1.7 | 0.9 ± 0.5 | 1.0 ± 1.3 | 0.9 ± 0.4 |
| 22:1n-9 | 2.2 ± 0.5 | 2.4 ± 0.4 | 3.3 ± 0.4 | 3.1 ± 0.4 |
| 22:1n-7 | 1.2 ± 0.6 | 1.0 ± 0.5 | 1.1 ± 0.3 | 1.6 ± 0.4 |
| 22:2n-6 | 1.4 ± 0.5 | 1.7 ± 0.6 | 3.0 ± 0.5 | 4.0 ± 0.8 |
| 21:5n-3 | 0.9 ± 0.6 | 1.8 ± 0.6 | 2.2 ± 0.6 | 1.6 ± 0.9 |
| 22:4n-6 | 0.2 ± 0.5 | 1.0 ± 1.6 | 0.3 ± 0.6 | 1.6 ± 1.7 |
| 22:5n-6 | 7.6 ± 1.1 | 10.7 ± 1.4 | 7.7 ± 0.7 | 9.6 ± 1.4 |
| 22:4n-3 | 2.3 ± 0.9 | 4.2 ± 1.3 | 5.1 ± 0.9 | 7.2 ± 1.7 |
| 22:5n-3 | 10.4 ± 0.9 | 10.8 ± 0.6 | 10.4 ± 2.4 | 11.1 ± 0.7 |
| 22:6n-3 | 33.9 ± 5.6 | 38.9 ± 4.3 | 23.1 ± 3.7 | 26.3 ± 4.7 |

All fatty acid values were converted to a mass percentage of the total array, and were named according the IUPAC nomenclature as X:Y n-z, where X is the number of carbon atoms in the fatty acids, Y is the number of methylene-interrupted double bonds in the chain, and n-z denotes the position of the last double bond relative to the methyl terminus (Ronconi et al., 2010). Fatty acids suggested by Iverson et al. (2004) as important dietary fatty acids, which transfer from prey to predator, were used in our analyses. Fatty acids profiles (% of fatty acids) were transformed using arcsin square-root function. Fatty acids groups were identified using a multivariate analysis R Package (Team, 2017), FactoMineR, using a hierarchical clustering analysis based on principal components (Husson et al., 2012). To reduce the number of variables used, A SIMPER (similarity percentage routine) was used to assess which fatty acids were primarily responsible for observed differences among groups (King et al., 1999). A principal components analysis (PCA) was performed on the fatty acid profiles with PC-ORD version 6 (McCune et al., 2011) among piscivorous groups to provide inferences about patterns of resource use as defined by Chavarie et al. (2016b). Two-way Permutational Multivariate Analysis of Variance (PERMANOVA), a non-parametric analog of Multivariate analysis of variance (MANOVA), was used to test for differences in fatty acid composition among the groups identified by FactoMineR and among arms (i.e., to investigate any spatial variations within the piscivorous ecotype). Two-way PERMANOVA were performed in PAST 3 (Hammer et al., 2001) using 9999 permutations. Pairwise *post-hoc* comparison (Bonferroni corrected) followed to test differences among groups defined by FactoMineR and among arms. Pairwise *post-hoc* comparison (Bonferroni corrected) also followed to test differences among arms (i.e., spatial variation). Finally, the fatty acid groups determined by FactoMineR were tested for differences in depth of capture using one-way analysis of similarities (ANOSIM) with 9999 permutations using PAST 3.

### Life-history

To determine if fatty acid groups differed in size-at-age, length vs. age was modeled using the Von Bertalanffy length-age model fit to length at age-of-capture of individual fish (Quinn et al., 1999):

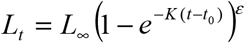

The length-age model describes length *L_t_* at age-of-capture *t* as a function of theoretical maximum length (*L_∞_* = mm), instantaneous rate at which *L_t_* approaches *L_∞_ (K* = 1/year), theoretical age-at-zero length (*t*_0_ = years), and multiplicative error (ε). Model parameters, *L*_∞_, *K*, and *t*_0_, and associated standard errors were estimated using nonlinear regression. Residual sums-of-squares were compared between a full model (separate models for each group) to a reduced model (a single model for all groups) in a likelihood-ratio test (Hosmer Jr et al., 2000). If the likelihood-ratio test was significant *(P* ≤ 0.05), we concluded that growth differed among groups identified by fatty acids. If the likelihood-ratio test was not significant (*P* > 0.05), we concluded that growth did not differ among groups. The same test was repeated for each pair of groups, with and without giant individuals (fork length ≥900 mm) included in each group, to isolate the influence of this subset in our size-at-age comparison due to the prevalence of giants in one of the groups (see Results).

### Genetic analyses

To determine if genetic differences existed among individuals expressing different feeding strategies, the 79 lake trout classified by fatty acid composition into four groups were genotyped to determine genetic variation and structure within and among groups. To allow a sample size sufficient for making a genetic comparison of giants to the other dietary groups, 22 additional individuals determined non-randomly by their size (≥ 900 mm; giant sub-set) from the 2002-2015 collections were added to giants processed for fatty acids, for a total of 39 giants. Lake trout DNA was extracted from pectoral fin tissue preserved in ethanol using DNEasy extraction kits (Qiagen Inc., Valencia, CA) following manufacturer protocols. Piscivorous groups were assayed using a suite of 23 putatively neutral microsatellite markers amplified in four multiplexes previously described in Harris et al. (2015). Amplified microsatellite fragments were analyzed using an automated sequencer (ABI 3130xl Genetic Analyzer; Applied Biosystems, Foster City, CA). The LIZ 600 size standard was incorporated for allele base-size determination. All genotypes were scored using GeneMapper software ver. 4.0 (Applied Biosystems) and then manually inspected to ensure accuracy.

The program MICROCHECKER ver. 2.2.0.3 (Van Oosterhout et al., 2004) was used to identify genotyping errors, specifically null alleles and large allele dropout. Observed and expected heterozygosity (H_E_ and *H_O_*) were calculated using GENEPOP ver. 4.2 (Rousset, 2008). The program HP-RARE ver. 1.1 (Kalinowski, 2005) was used to determine the number of alleles, allelic richness, and private allelic richness for each group, sampling 22 genes in each sample. Tests of departure from Hardy-Weinberg equilibrium and genotypic linkage disequilibrium within each sample (i.e., for each fatty acid grouping and the Giant subset) were conducted in GENEPOP using default values for both. Results from all tests were compared with an adjusted alpha (α = 0.05) following the False Discovery Rate procedure (Narum, 2006).

We used the POWSIM V. 4.1 analysis to assess the statistical power of our microsatellite data set given the observed allelic frequencies within our samples in detecting significant genetic differentiation between sampling groups (Ryman et al., 2006). For POWSIM analyses, we assumed that lake trout within our study diverged from a common baseline population with the same allelic frequencies as observed in our contemporary samples. Simulations were performed with an effective population size of 5000 to yield values of F_ST_ of 0.01, 0.005 and 0.001. The significance of tests in POWSIM were evaluated using Fisher’s exact test and the χ2 test and the statistical power was determined as the proportion of simulations for which these tests showed a significant deviation from zero. All simulations were performed with 1000 iterations.

Genetic structuring was tested among lake trout groups using several different methods. First, genotypic differentiation among lake trout groups was calculated using log-likelihood (G) based exact tests (Goudet et al., 1996) implemented in GENEPOP. Global F_ST_ (θ) (Weir et al., 1984) was calculated in FSTAT ver. 2.9.3 (Goudet, 1995) and pairwise comparisons of F_ST_ between groups were calculated in ARLEQUIN ver. 3.5 (Excoffier et al., 2005) using 10,000 permutations. We then employed the Bayesian clustering program STRUCTURE V. 2.3.2 (Pritchard et al., 2000) to resolve the putative number of populations (i.e., genetic clusters (K)) within our samples. Owing to the remarkably low levels of genetic differentiation among lake trout in the Great Bear Lake (Harris et al., 2015; Harris et al., 2013), we employed the LOCPRIOR algorithm (Hubisz et al., 2009). The LOCPRIOR algorithm considered the location/sampling information as a prior in the model, which may perform better than the traditional STRUCTURE model when the genetic structure is weak (Hubisz et al., 2009). We also incorporated an admixture model with correlated allelic frequencies and the model was run with a burn-in period of 500,000 iterations and 500,000 Markov chain Monte Carlo iterations. We varied the potential number of populations (K) from 1 to 10 and we ran 20 iterations for each value of K. The STUCTURE output was first processed in the program STRUCTURE HARVESTER (Earl, 2012), followed by the combination of results of independent runs of the program and compilation of results based on lnP(D) and the post hoc ΔK statistic of Evanno et al. (2005), to infer the most likely number of clusters. The best alignment of replicate runs was assessed with CLUMPP V. 1.1 (Jakobsson et al., 2007) and DISTRUCT V. 1.1 (Rosenberg, 2004) was then used to visualize the results. For STRUCTURE analyses, we reported both lnP(D) and the post hoc ΔK statistic.

Finally, Discriminant Analysis of Principal Components (DAPC) (Jombart et al., 2010) was implemented in the Adegenet package (Jombart, 2008) in R (Team, 2015). The number of clusters was identified using the *find.clusters* function (a sequential K-means clustering algorithm) and subsequent Bayesian Information Criterion (BIC), as suggested by Jombart et al. (2010). Stratified cross-validation (carried out with the function *xvalDapc)* was used to determine the optimal number of principal components to retain in the analysis.

### Morphology

Morphological variation was quantified for the 79 lake trout and used to compare fatty acid groupings (different feeding strategies) identified within the piscivorous ecotype. Twenty-three landmarks and 20 semi-landmarks, based on Chavarie et al. (2015), and fourteen linear measurements based on Muir et al. (2014), were used to characterize body and head shape from digital images. The combination of traditional and geometric ecotype metrics was used because relationships of phenotype morphology with foraging (e.g., jaw size) and swimming (e.g., fin lengths and caudal peduncle depth) (Kahilainen et al., 2004; Kristjánsson et al., 2002; Webb, 1984). Landmarks and semi-landmarks were digitized in x and y coordinates using TPSDig2 software (http://life.bio.sunysb.edu/ecotype). Subsequently, digitized landmarks and semi-landmarks were processed in a series of Integrated Morphometrics Programs (IMP) version 8 (http://www2.canisius.edu/;sheets/ecotypesoft), using partial warp scores, which are thin-plate spline coefficients. Morphological methods and programs are described in Zelditch et al. (2012) and specific procedures were described in further detail by Chavarie et al. (2013). All morphological measurements were size-free, using centroid sizes or residuals from regressions on standard length (Zelditch et al., 2012).

Canonical Variate Analyses (CVA) were conducted on all morphological data, including body shape, head shape, and linear measurements, to determine relationships among fatty acid groups. Body and head shape were analysed using CVAGen8 from the IMP software (Zelditch et al., 2012) and for linear measurements, CVA was analyzed with SYSTAT (Systat Software Inc., Chicago, IL, USA). Single Factor Permutation MANOVA with 10 000 permutations tested for differences among groups and determined the percentage of variation explained for a grouping if a CVA was significant. For linear measurements, a Bonferroni-corrected post-hoc test followed MANOVA to identify measurements that differed among group. Principal component analyses (PCA) were performed on body- and head-shape data using PCAGen8 (IMP software) among groups to visualize morphological variation within the dataset. PC-ORD version 6 software (McCune et al., 2011) was used to perform a PCA on the linear measurements.

## Results

### Fatty acids

On the basis of fatty acid composition, piscivorous lake trout were divided along a resource use axis into four groups (1-4; Fig. A2), containing 14, 16, 21, and 28 individuals, respectively (Figs. 2 and A2; Table 1). Average dissimilarity was 14.61 (SIMPER analysis); whereas, the most discriminating 26 fatty acids, explaining together ~89% of the separation among groups, were: 22:6n-3 (12.5 %), 18:1n-9 (10.8 %), 16:1n-7 (6.8 %), 20:5n-3 (5.0 %), 20:4n-6 (3.9 %), 18:2n-6 (3.8 %), 22:4n-3 (3.7 %), 16:0 (3.5%), 20:4n-3 (3.3%), 18:1n7 (3.3%), 20:2n-6 (3.1%), 14:0 (2.8%), 20:1n-9 (2.7%), 22:5n-6 (2.7%), 20:3n-3 (2.3%), 22:2n-6 (2.1%), 18:0 (2.0%), 18:3n-3 (1.9%), 18:4n-3 (1.8%), 22:4n-6 (1.7%), 20:1n-7 (1.5%), 22:5n-3 (1.4%), 21:5n-3 (1.3%), 22:1n-11 (1.2%), 20:0 (1.2%), 16:4n-3 (1.2%), and 16:2n-4 (1.1%) (Table 1). The first two axes of the fatty acids PCA explained 65.2 % of the variation in diet and the four groups were supported by PERMANOVA (F_3,76_ = 23.9, P < 0.01) and pairwise comparisons between all pairs (all P < 0.01; Bonferroni corrected). Spatial differences in fatty acids composition were found among arms (F_4,76_ = 3.2, P < 0.01). Pairwise comparisons identified differences between Smith and McVicar arms (P = 0.02; Bonferroni corrected; Fig. A3). Interaction between fatty acids groups and arms was not significant (p > 0.05). Finally, depth of capture did not differ among fatty acid groups (p > 0.05). For all groups, most lake trout were caught between 0 and 20 m (Fig. A4).

**Fig. 2.**
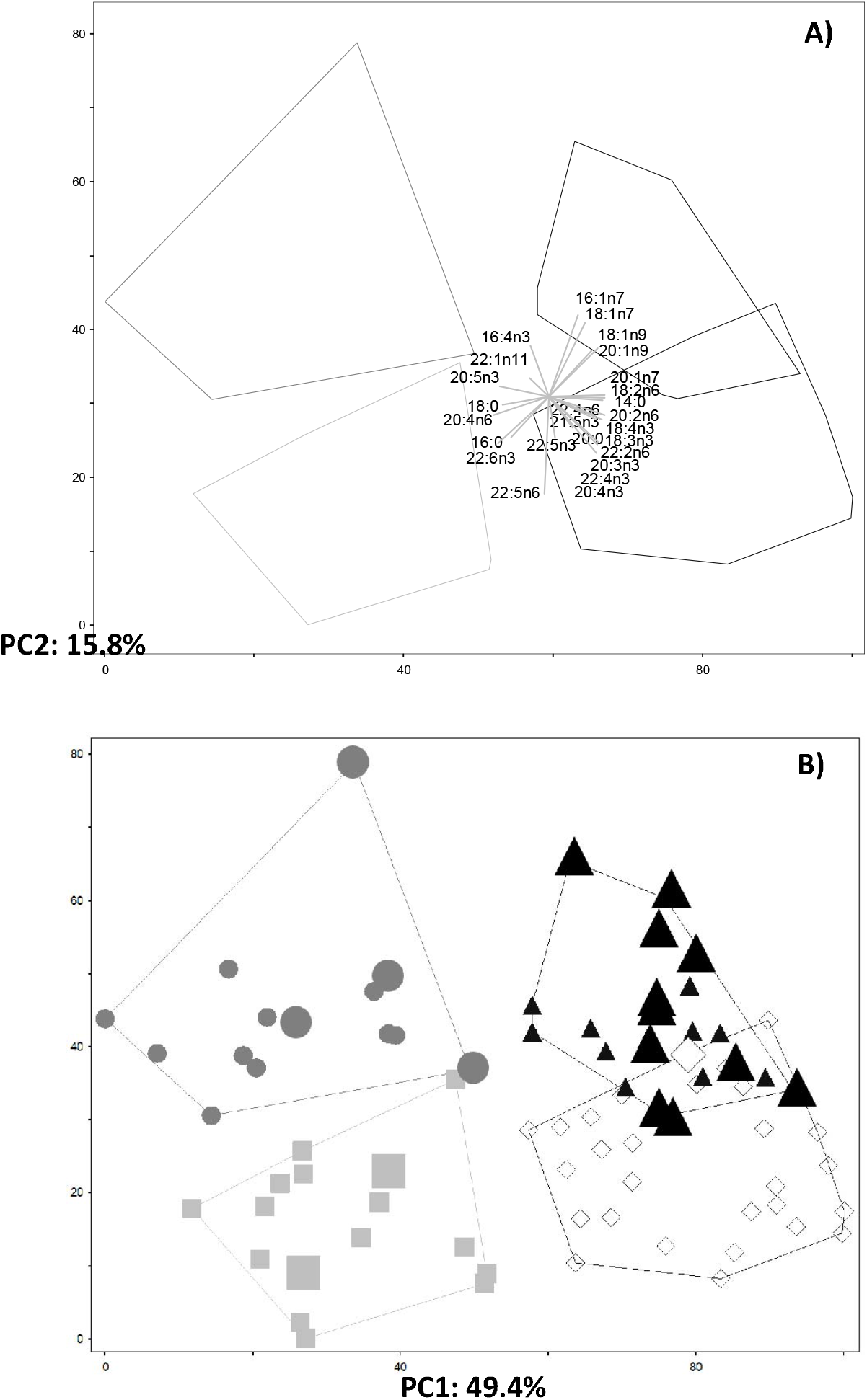
Principal Components Analysis of fatty acids of 79 Lake Trout classified as the piscivorous morph from Great Bear Lake, based on the most discriminating 26 fatty acids from SIMPER analysis, explaining together ~89% of the separation among groups. A) Vectors of individual fatty acids contributing to the positioning of piscivorous individuals and the convex hull delimitating group’s position are shown. B) Individual Lake Trout are represented as circle = Group 1, square = Group 2, triangle = Group 3, and diamond = Group 4. To visualize their variation within and among groups, large symbols were used to depict individuals longer than 900 mm fork length, which were identified as the Giant sub-set in this study. Groups were defined by FactoMineR using fatty acids and they are outlined by convex hulls.

### Life-history

Overall, life history parameters did not differ among lake trout groups grouped by fatty acid composition, including length-age models (Fig. 3; *F*_9, 63_ = 1.58; *P* = 0.141). With the giant sub-set included, growth differed between only between Group 3 and Group 4 (*F*_3, 41_ = 3.958; *P* = 0.014), but not between any other pairs (P > 0.1). Without Giants included (prevalence of Giants was higher in Group 3 than Group 1, Group 2, and Group 4), none of the pairs differed for length-at-age (P > 0.1), suggesting growth rate similarities among groups.

**Fig. 3.**
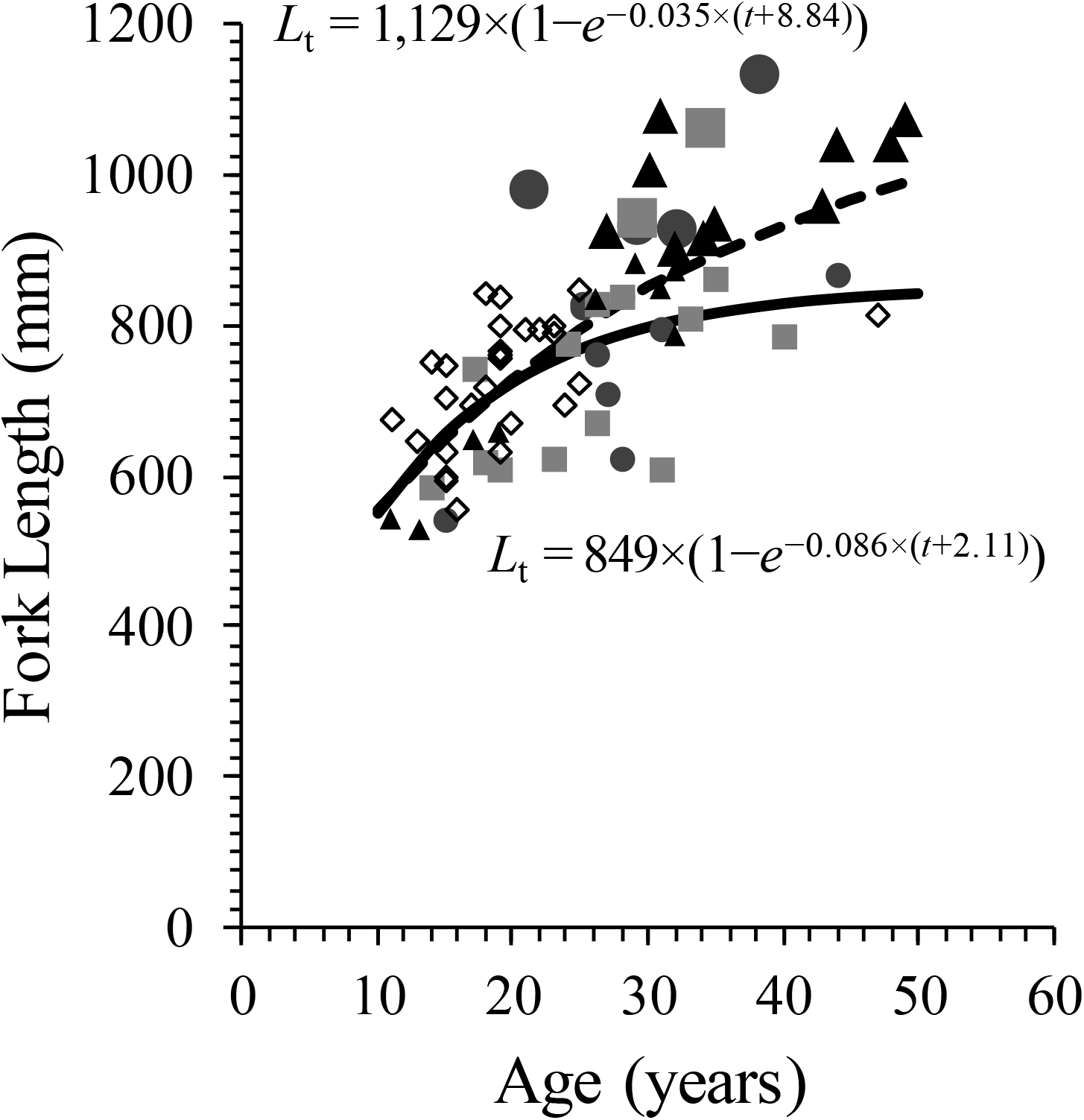
Fork length (mm) at age (years) for four groups of piscivorous Lake Trout sampled from Great Bear Lake in 2002–2015 (Group 1 = squares; Group 2 = circles; Group 3 = triangles; diamond = Group 4). Large symbols depict Giants (FL > 900 mm) within each group. The von Bertalanffy length-age models are depicted as a solid line (without Giants) and a dashed line (with Giants).

### Genetic differentiation

Piscivorous lake trout groups displayed little genetic differentiation, except for the Giant sub-set, which differed slightly from other groups that were defined by fatty acids. MICROCHECKER identified two loci (OtsG253b and Sco102) that contained null alleles. These loci, along with non-variable loci Sco218 and SSOSL456, were removed, leaving 19 informative loci for subsequent analyses. Descriptive statistics of genetic variation were similar among groups. The number of alleles per locus ranged from four (Smm21) to 41 (SnaMSU10) and averaged 28.75 across all loci. Averaged observed heterozygosity ranged from 0.78 (Giant) to 0.83 (Group 1) while expected heterozygosity was 0.84 for all groups except Group 1 (0.85; Table 2). Allelic richness ranged from 9.57 (Group 2 and 4) to 9.87 (Group 1), while expected private allelic richness ranged from 0.87 (Group 3) to 1.08 (Group 2; Table 2). Only five of 95 tests (all of which involved different loci) showed significant departures from Hardy-Weinberg equilibrium after adjustment for False Discovery Rate (adjusted α = 0.01). Of those five, all were heterozygote deficits and three involved the Giant sub-set. Only nine of 885 tests revealed significant linkage disequilibrium after adjusting for False Discovery Rate (adjusted α = 0.0068). No locus-pair linkage disequilibrium combinations were consistently significant, but seven of nine departures were in the Giant sub-set.

**Table 2.** Number of individuals genotyped (*N*), number of alleles (*N_A_*), expected heterozygosity (*H*_E_), observed heterozygosity (*H*_O_), allelic richness (*A*_R_) and private allelic richness (*PA*_R_) within fatty acid groups identified within a piscivorous morphotype of Lake Trout from Canada’s Great Bear Lake.

|  | **N** | **N<sub>A</sub>** | **H<sub>E</sub>** | **H<sub>O</sub>** | **A<sub>R</sub>** | **PA<sub>R</sub>** |
| --- | --- | --- | --- | --- | --- | --- |
| **Group 1** | 12 | 10.16 | 0.85 | 0.83 | 9.87 | 1.08 |
| **Group 2** | 16 | 11.26 | 0.84 | 0.82 | 9.57 | 0.99 |
| **Group 3** | 20 | 12.32 | 0.84 | 0.81 | 9.70 | 0.87 |
| **Group 4** | 28 | 14.11 | 0.84 | 0.81 | 9.57 | 0.98 |
| **Giant** | 39 | 15.95 | 0.84 | 0.78 | 9.69 | 1.05 |

Using our microsatellite data set, the POWSIM analysis indicated a 100% power of detecting F_ST_ values as low of 0.01 and 0.005. However, power was reduced to 77% when assessing genetic differentiation at a F_ST_ of 0.001. Overall, our microsatellite data set (including the number of loci, alleles per locus, and sample sizes) had sufficient power to detect relatively low levels of genetic differentiation.

Global genetic differentiation was extremely low (θ = 0.001, 95% c.i. = −0.002-0.005) among the groups of piscivorous lake trout. Pairwise F_ST_ ranged from −0.004 to 0.016 (Table 3); comparisons that included Giants always differed the most from the other fatty acid groups, and they were involved in the only significant pairwise comparisons (P < 0.05, Table 3). The F_ST_ values for the Giant vs. Groups 1 and 4 were generally similar to genetic differentiation among the four original lake trout ecotypes in Great Bear Lake, except for Ecotype 1 vs Ecotype 2 (Table 3). Bayesian clustering implemented in STRUCTURE provided evidence for two genetic clusters when evaluating both lnP(D) or ΔK (Table A2). The admixture plot based on K=2 showed no clear genetic structure between groups defined by fatty acid analysis; however, some differentiation of the Giant sub-set from the fatty acid groups was observed (Fig. 4).

**Fig. 4.**
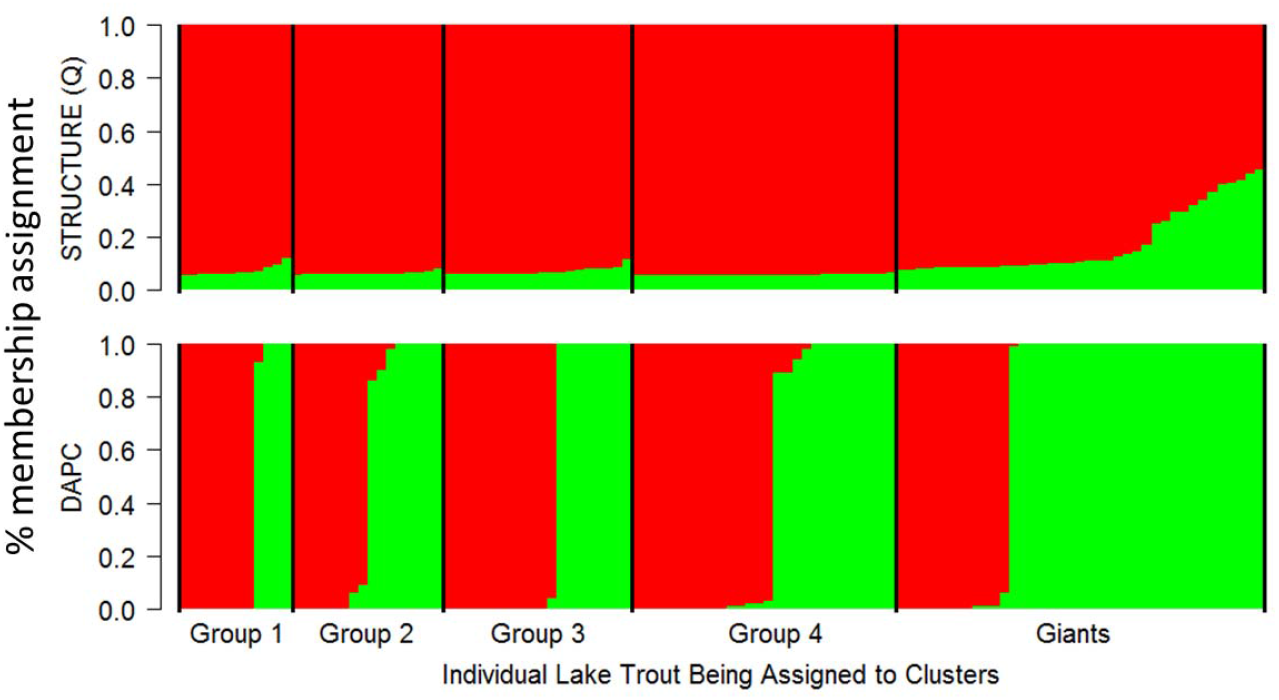
Results of the Bayesian clustering analysis implemented in the program STRUCTURE (B) and the compoplot of percent membership assignment revealed from the DAPC analysis (B) for piscivorous Lake Trout from Great Bear Lake. Shown is the admixture coefficient/percent membership assignment plot where each individual is represented as a vertical line partitioned into colored segments representative of an individual’s fractional membership in any given cluster (K). The most likely number of genetic clusters was two in both the STRUCTURE analysis (based on lnP[D] and the ΔK statistic of Evanno et al. (2005)) and DAPC analysis (based on the lowest BIC score and with 30 PCs retained).

**Table 3.** Pairwise F_ST_ based on variation at microsatellite loci among Lake Trout morphs from Harris et al. (2015) and piscivorous fatty acids dietary groups from Great Bear Lake. Significant results are represented as follow: * values are significant at an initial α of 0.05 and ** values are significant at an α of 0.02 subsequent False Discovery Rate adjustments for multiple comparisons.

|  | Morph 1 | Morph 2 | Morph 3 |  | Group 1 | Group 2 | Group 3 | Group 4 |
| --- | --- | --- | --- | --- | --- | --- | --- | --- |
| Morph 1 |  |  |  | Group 1 |  |  |  |  |
| Morph 2 | 0.063** |  |  | Group 2 | 0.003 |  |  |  |
| Morph 3 | 0.004** | 0.007** |  | Group 3 | 0.001 | -0.01 |  |  |
| Morph 4 | 0.012** | 0.017** | 0.009** | Group 4 | 0.005 | -0.004 | -0.002 |  |
|  |  |  |  | Giant | 0.016** | 0.001 | -0.002 | 0.006** |

Finally, the Bayesian information criterion in the DAPC analysis (BIC = 185.42, Table A3, Fig. A5 A) suggested that two clusters best explained genetic structure in our study (30 PCs retained as suggested by the cross-validation procedure; Fig. A5 B). A compoplot (barplot showing the probabilities of assignment of individuals to the different clusters) for K=2 revealed no clear genetic structure between two groups identified by the DAPC analysis except for the Giant group, which appeared to have more individuals assigned to cluster two (Fig. 4). Density plots of the discriminant function, however, suggested that the two clusters identified through the DAPC analysis are mostly non-overlapping (Fig. A5 C).

### Morphology

Morphological variation was low among the four dietary groups within the piscivorous ecotype. The first canonical axis for body shape CVA was significant (P > 0.05), but head shape CVA revealed no significant canonical axes (P > 0.05) in groupings (Fig. 5 a, b, c). MANOVAs for body and head shape were not significant (P > 0.05). Linear measurements CVA revealed one significant canonical axis (P > 0.05). MANOVA permutation tests confirmed differences in linear measurements among groups (P = 0.047). Most distinctions were related to linear measurements of heads, with upper and lower jaws, head depth, and snout-eye lengths differing between Group 3 and Group 4 (P ≤ 0.05), and head length differing between Group 1 and 4 (P = 0.03; Fig. 6). Caudal peduncle length and anal fin length differed marginally between Groups 2 vs 3 (P = 0.068) and Groups 1 vs 3 (P = 0.075), respectively. The first two PCA axes explained 44.3% and 12.3 % of variation for body shape, 35.1% and 30.7 % of variation for head shape, and 39.6 % and 20.9 % for linear measurements (Fig. 5 d, e, f).

**Fig. 5.**
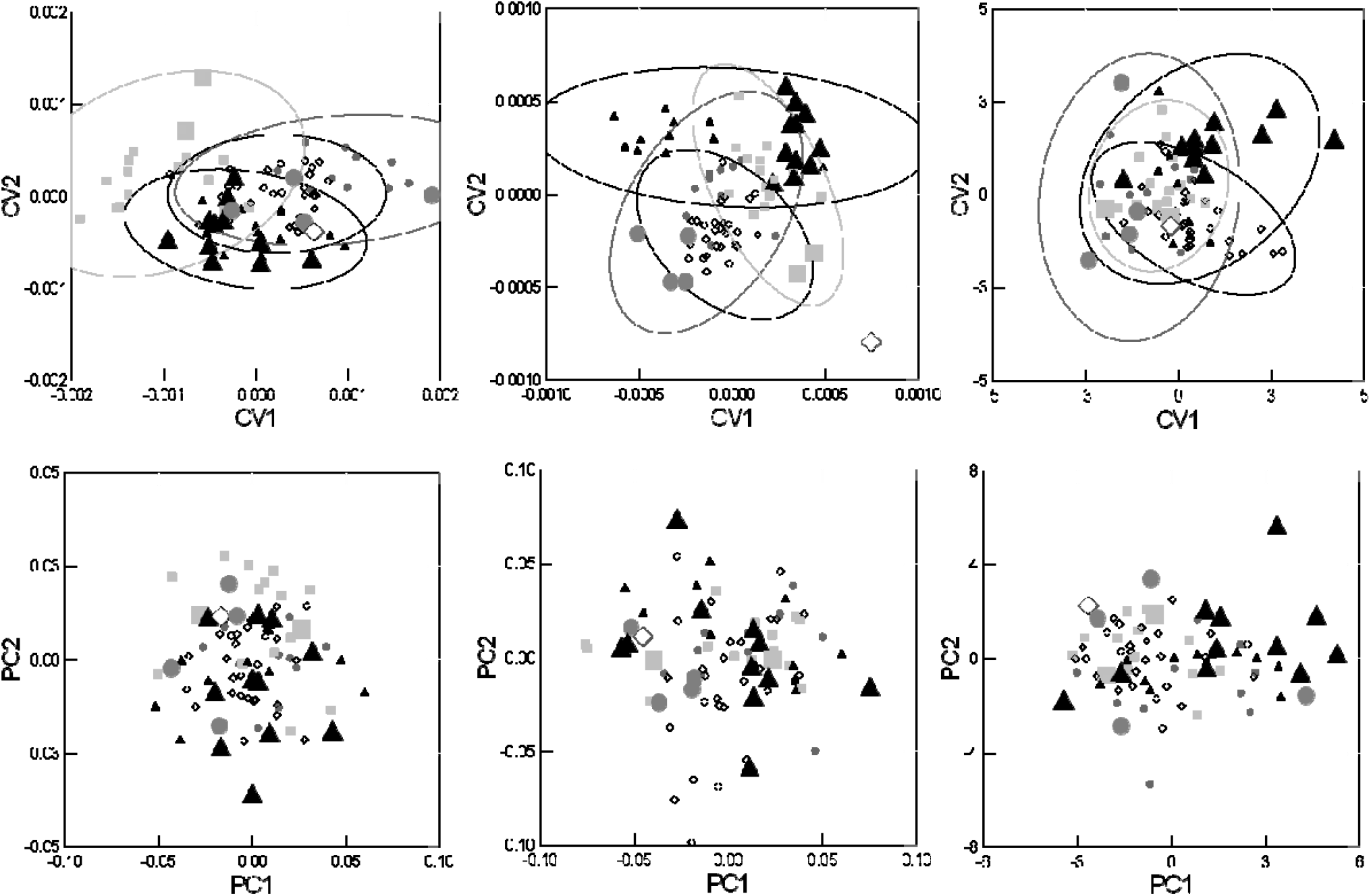
Canonical Variate Analyses (95% ellipses) and Principal Components Analysis of body shape (a, d), head shape (b, e) and linear measurements (c, f), respectively, of piscivorous Lake Trout represented as: square = Group 1, circle = Group 2, triangle = Group 3, and diamond = Group 4. The first two PCA axes explained 44.3% and 12.3 % of variation for body shape, 35.1% and 30.7 % of variation for head shape, and 39.6 % and 20.9 % for linear measurements (Fig. 6 d, e, f). To visualize their variation within and among groups, individuals longer than 900 mm FL, which are considered the Giant sub-set in this study, are depicted by larger symbols.

**Fig. 6.**
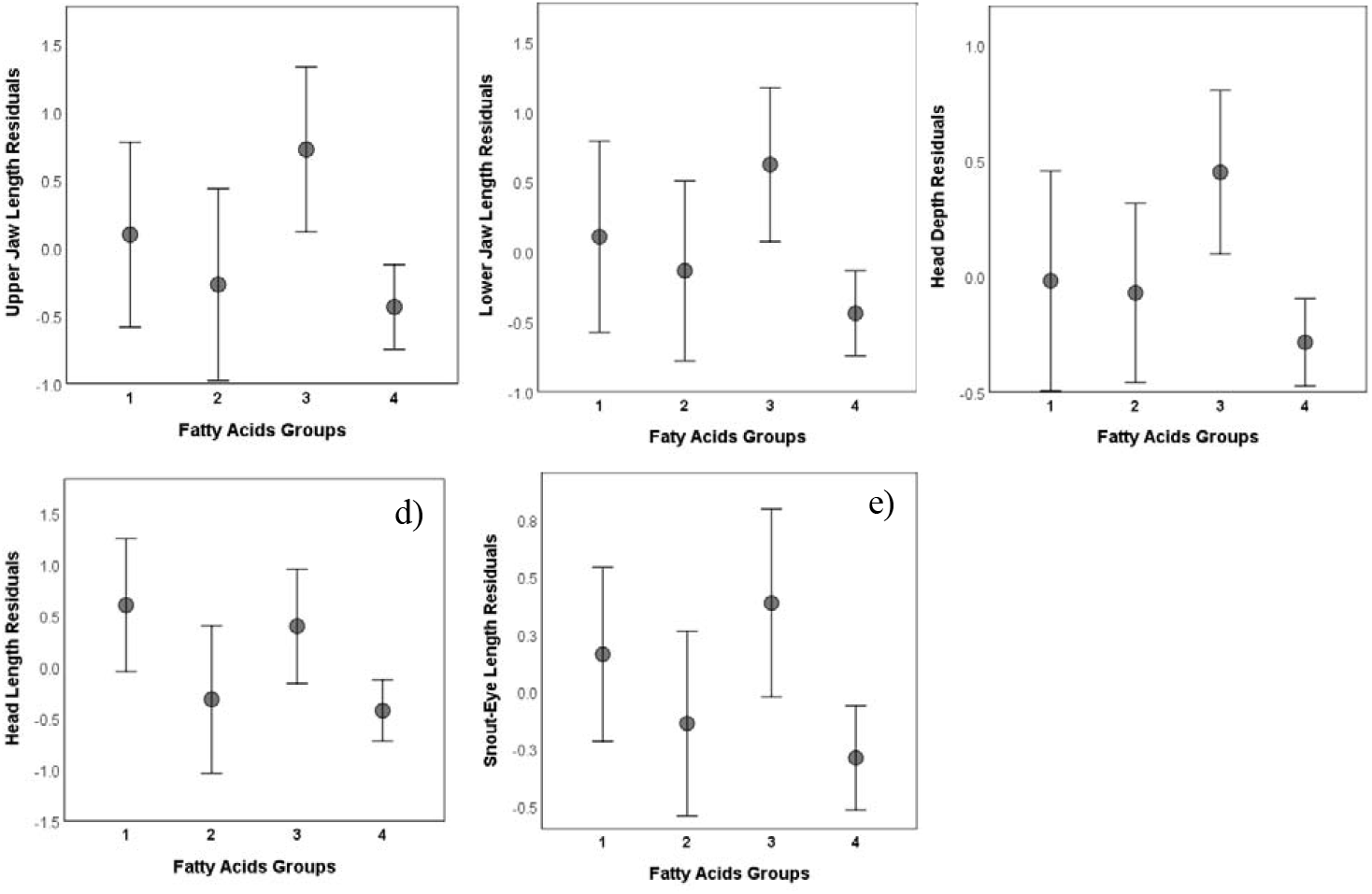
Residuals of mean (± 95%CI) size-standardized upper and lower jaw lengths, head depth and length, and snout-eye length among piscivorous Lake Trout groups. Grouping symbols are as follows: square = Group 1, circle = Group 2, triangle = Group 3, and diamond = Group 4.

## Discussion

A common assumption in polyphenism is that partitioning and variability of resource use will occur predominantly among ecotypes rather than within ecotypes. In contrast, homogeneity of resource use is anticipated to occur within ecotypes, be spatially and temporally stable, and provide the selection opportunity for specialization (Amundsen et al., 2008; Knudsen et al., 2010; Svanbäck et al., 2004). However, this study provided evidence that variation occurred within an ecotype due to diet specialization among individuals, possibly a precursor to further population diversification via fine scale ecological selection (Richardson et al., 2014; Vonlanthen et al., 2009). The co-existence of multiple generalist ecotypes in Great Bear Lake (Chavarie et al., 2016a), combined with the individual specialization shown here in the piscivorous generalist ecotype, expands our understanding of niche use and expansion, plasticity, individual specialization, and intraspecific diversity in evolutionarily young populations.

Using fatty acids as dietary biomarkers, four distinct patterns of resource use were identified within the piscivorous lake trout of Great Bear Lake (Fig. 2). Groups 3 and 4 had the most overlap and these groups were characterized by C20 and C22 monounsaturates, biomarkers of a food web based on pelagic or deep-water copepods (Ahlgren et al., 2009; Happel et al., 2017; Hoffmann, 2017; Loseto et al., 2009; Stowasser et al., 2006). Specifically, 20:1n-9 is associated with calanoid copepods known to be particularly important in northern pelagic food webs (Ahlgren et al., 2009; Budge et al., 2006; Kattner et al., 1998; Loseto et al., 2009). High levels of 14:0, 18:3n-3 and 18:4n-3 fatty acids within groups 3 and 4 are also associated with pelagic environments (Scharnweber et al., 2016; Tucker et al., 2008), although high levels of 18:2n-6 and 18:3n-3 have also been associated with terrestrial markers (Budge et al., 2001; Budge et al., 1998; Hoffmann, 2017).

Groups 1 and 2 were characterized by high concentrations of 16:4n-3, 20:4n-6 and 22:6n-3 found in diatom and dinoflagellate-based food webs, respectively. The fatty acid 20:4n-6 reflects a benthic feeding strategy (from benthic invertebrates to fish) (Stowasser et al., 2006; Tucker et al., 2008), whereas 22:6n-3 in pennate diatoms (Iverson, 2009) and filter feeders links planktonic dinoflagellates to benthic filter-feeding bivalves in a food web (Alfaro et al., 2006; Virtue et al., 2000). Relatively high concentrations of 16:0, 18:0 and 22:6n-3 and low concentrations of 16:1n-7 supported the interpretation of carnivorous (or cannibalistic) dietary patterns (Dalsgaard et al., 2003; Iverson, 2009; Iverson et al., 2004; Piché et al., 2010). Individuals positioned between ends of principal components suggests a clinal pattern of resource use or habitat coupling (Vonlanthen et al., 2009), where borders among groups are neither abrupt nor obvious as they are part of a continuum (Hendry et al., 2009b). Overall, observed trophic patterns could reflect prey associated with different microhabitat patches; however, the key assumption of disparity of prey associated with habitat heterogeneity (Skulason et al., 1995; Svanbäck et al., 2005) may not be applicable to Great Bear Lake (Chavarie et al., 2016a; Chavarie et al., 2020).

Sympatric divergence, in which barriers to gene flow are driven by selection between ecological niches, has been implicated in the evolution of ecological and morphological variation in fishes (Chavarie et al., 2016d; Hendry et al., 2007; Præbel et al., 2013). Despite the limited ability of neutral microsatellite markers to detect patterns of functional divergence (Berg et al., 2016; Lamichhaney et al., 2016; Roesti et al., 2015), the significant genetic differentiation based on comparisons with Giant sub-set suggests some deviation from panmixis within the piscivorous ecotype. Such a genetic pattern displayed by the Giant sub-set, despite a lack of ecological discreteness, perhaps resulted from size-assortative mating and/or differences in timing and location of spawning (Nagel et al., 1998; Rueger et al., 2016; Servedio et al., 2011). Great Bear Lake is not the only lake in North America with an apparent divergence in lake trout body size; in Lake Mistassini, “Giant” individuals also differed genetically from other lake trout groups (Marin et al., 2016). The similarity based on lake trout body size between both lakes suggests analogous variables favoring partial reproductive isolation. Although alternative causes of genetic differentiation may be possible, due to the short time since the onset of divergence, post-zygotic isolation seems unlikely in this system (e.g., prezygotic isolation generally evolves more rapidly Coyne et al., 2004) and we therefore favor assortative mating based on size and location as an explanation for the low-level genetic divergence observed. Nonetheless, putative partial reproductive isolation within an ecotype adds to the complexity of diversification and speciation processes potentially occurring within lake trout in Great Bear Lake (Hendry, 2009; Nosil et al., 2009).

A central question arising from our analysis is what are the mechanisms behind these patterns of variation? As individual specialization can result in dietary sub-groups and perhaps differences in habitat use among sections of a population, such inter-individual variation within ecological sub-groups could substantially influence processes of diversification (Araújo et al., 2008; Cloyed et al., 2016). Among-individual resource specialization within an ecotype in a species-poor ecosystem like Great Bear Lake could reflect the diversifying force of intraspecific competition, lack of constraining effects of interspecific competition, the abundance and distribution of resources (e.g., temporal and spatial variation of resources), or some combination of these variables (Bolnick et al., 2007; Cloyed et al., 2016). Multiple patterns of resource specialization within a single ecotype, as we see for lake trout in Great Bear Lake, contrasts with the expected pattern of trophic divergence among ecotypes and homogenization in habitat use or diet within an ecotype, a key assumption guiding the development of functional ecological theory (Svanbäck et al., 2004; Violle et al., 2012). Expression of intraspecific divergence through habitat and foraging specialization is thought to drive selection on traits that enable more efficient use of resources (Schluter, 2000; Skulason et al., 1995; Snorrason et al., 2004).

In Great Bear Lake, multiple trophic generalists (which include the piscivores studied herein) coexist with one specialist lake trout ecotype. This contrasts with the more commonly reported observation of multiple specialist ecotypes (Chavarie et al., 2016a; Elmer, 2016; Kassen, 2002). A generalist population, however, can be composed of several subsets of specialized individuals (Bolnick et al., 2009; Bolnick et al., 2007; Bolnick et al., 2002). This broad distribution of trophic variation within a population appears to be the case within the Great Bear Lake piscivores. The among-individual specialization may result, to some degree, from variable use of spatially separated resources and possibly temporally variable resources, both of which could be expected in a large northern lake (Fig. A4; Costa et al., 2008; Cusa et al., 2019; Quevedo et al., 2009). Ecologically, among-individual resource specialization within an ecotype is another form of diversity (Araújo et al., 2008; Bolnick et al., 2003; Pires et al., 2011). Such diversity may increase stability and persistence of an ecotype within a system where energy resources are scarce and ephemeral (Cloyed et al., 2016; Pfennig et al., 2012; Smith et al., 2011). Whether the level of among-individual specialization within this ecotype is stable or not is a question that cannot be answered with our data.

Realized niche expansions are often linked to individuals of different morphologies and body sizes, with evidence of efficiency trade-offs among different resources (Cloyed et al., 2016; Parent et al., 2014; Roughgarden, 1972; Svanbäck et al., 2004). When a resource gradient exists, niche expansion can be achieved via genetic differentiation, phenotypic plasticity, or a combination of these processes (Bolnick et al., 2020; Parent et al., 2014). The apparent segregation of resource use, based on our fatty acid analyses, despite a lack of major morphological, body size, and genetic differentiation among the four dietary groups within the piscivorous ecotype, suggests that behavioral plasticity is causing the observed patterns of dietary differentiation. Plasticity may promote diversification by expanding the range of phenotypes on which selection can act (Nonaka et al., 2015; Pfennig et al., 2010; West-Eberhard, 2003). Theoretical models suggest that exploiting a wide range of resources is either costly or limited by constraints, but plasticity is favored when 1) spatial and temporal variation of resources are important, 2) dispersal is high, 3) environmental cues are reliable, 4) genetic variation for plasticity is high and 5) cost/limits of plasticity are low (Ackermann et al., 2004; Hendry, 2016).

The expression of plasticity in response to ecological conditions (e.g., habitat structure, prey diversity) can increase fitness. While most studies of diet variation focus on morphological differences among ecotypes in a population, diet variation can also arise from behavioral, biochemical, cognitive, and social-rank differences that cause functional ecology to be expressed at a finer scale than at the ecotype level (McGill et al., 2006; Svanbäck et al., 2005; Violle et al., 2012; Zhao et al., 2014). Indeed, behavioral plasticity likely has a temporal evolutionary advantage due to relatively reduced reliance on ecologically beneficial morphological adaptation (Smith et al., 2011; Svanbäck et al., 2009). The only detectable morphological differences among piscivorous groups we identified in Great Bear Lake were associated with jaw lengths, snout-eye distance, and head length and depth, which are strongly related to foraging opportunities (Adams et al., 2002; Sušnik et al., 2006; Wainwright et al., 2016). Some morphological characters likely express different degrees of plastic responses (adaptive or not), and thus may be expressed differently depending on the magnitude and time of exposure to heterogeneous environments (Hendry, 2016; Sharpe et al., 2008). For example, environmental components (e.g., habitat structure) appear to have stronger and faster effects on linear characters (e.g., jaw length) than on body shape (Chavarie et al., 2015; Sharpe et al., 2008). Trophic level might also limit the scope for morphological variation in lake trout because piscivory can limit diversification of feeding morphology in fishes (Collar et al., 2009; Svanbäck et al., 2015).

## Conclusion

Understanding ecological mechanisms of diversification is challenging (Ackermann et al., 2004). Divergence occurs along a continuum and in early stages, such as in post-glacial lakes, morphological and dietary variation may not always be features that are related (Bolnick et al., 2020; Bolnick et al., 2007). The debate around diversification sequence, (which diverges first, behaviour, morphology, or ecology?) highlights the mosaic nature of intraspecific variation (Hendry et al., 2009a). In this study, we asked whether among-individual diet variation could be occurring within an ecotype by examining the fine-scale trophic variation of an early stage of sympatric divergence of lake trout in Great Bear Lake (i.e., postglacial, representing ~350 generations; Harris et al., 2015). Due to presumed homogeneity, few studies have investigated dietary patterns and groupings within an ecotype. Thus, this study provides evidence that among-individual resource specialization can occur within an ecotype.

Rapid divergence within relatively few generations and among-individual diet variation have both been demonstrated to be a strong driver of population dynamics (Ashley et al., 2003; Bolnick et al., 2020; Fussmann et al., 2007; Turcotte et al., 2011). In this study, the fine-grained trophic patterns shown within this ecotype suggested that ecological drivers (i.e., spatial variation, habitat use, prey diversity, and abundance) could have important effects on plasticity expression in early stages of divergence. Theory and experiments have demonstrated that among-individual diet variation can increase stability within a system (Agashe, 2009). Using a broad resource spectrum has been identified as an adaptive strategy for fishes living in Arctic environments, where food availability is patchily distributed and ephemeral (Dill, 1983; Kassen, 2002; Smith et al., 2011). Thus, it is no surprise that the trophic individual specialization within an ecotype was discovered within a northern lake.

## List of abbreviations

BP: before present
m: meter
mm: millimeter
h: hour
ca.: around
i.e.,: stands for
e.g.,: for example
NaCl: Sodium Chloride
FAME: fatty acid methyl esters
H_2_SO_4_: Sulfuric acid
GC: Gas chromatographic
°C: degree Celsius
°C/min: degree Celsius/minutes
UPGMA: Unweighted Pair Group Method with Arithmetic Mean
PCA: principal component analysis
PERMANOVA: Permutational Multivariate Analysis of Variance
MANOVA: Multivariate analysis of variance
SIMPER: similarity percentage routine
ANOSIM: analysis of similarities
*N*: Number of individuals genotyped
*N_A_*: number of alleles
*H*_E_: expected heterozygosity
*H*_O_: observed heterozygosity
*A*_R_: allelic richness
*PA*_R_: private allelic richness
A: alpha
FCA: Factorial correspondence analysis k=number of alleles
DAPC: Discriminant Analysis of Principal Components
IMP: Integrated Ecotypeometrics Programs
CVA: Canonical Variate Analyses

## Acknowledgements

We thank one anonymous reviewer and Alexandra Tyers for their insightful readings and constructive suggestions. Déline Renewable Resources Council, Déline Lands and Finance Corporation, the community of Déline, DFO in Hay River, and the Department of Environment and Natural Resources in Déline provided valuable help with field planning and logistics. We especially thank J. Chavarie, S. Buckley, L. Harris, G. Lafferty, M. Lindsey, M. Low, Z. Martin, S. Wiley, and C. Yukon, who helped lead sampling teams and coordinate logistics. The following individuals who helped conduct field sampling in various years are gratefully acknowledeged: J. Baptiste, D. Betsidea, D. Baton, L. Dueck, R. Eshenroder, G. Menacho, N. Modeste, I. Neyelle, L. Neyelle, M. Smirle, A. Swietzer, C. Takazo, A. Vital, F. Vital, B. Yukon, M. Yukon, T. Yukon and Charity, Cameron, and Cyre Yukon. D. Phillibert, K. Laubriat, S. Thacker, and K. Theoret conducted isotopic lab work.

## Declarations

### Authors’ contributions

LC, KH, WT, CK, and AM conceived and funded the study. LC and CG carried out the field work. LC, CG, LH, and MH participated in the data analyses. LC wrote the manuscript. All authors read and approved the final manuscript.

## Availability of data and materials

The datasets supporting the conclusions of this article are included within the article. Raw data are available from the corresponding author upon reasonable request.

## Consent to publish

Not applicable.

## Ethics approval and consent to participate

We declare that our experiments were performed in the respect of ethical rules. This protocol was approved by Department of Fisheries and Ocean Canada, Freshwater Institute Animal Care Committee Science Laboratories.

## Appendix

**Table A1.**
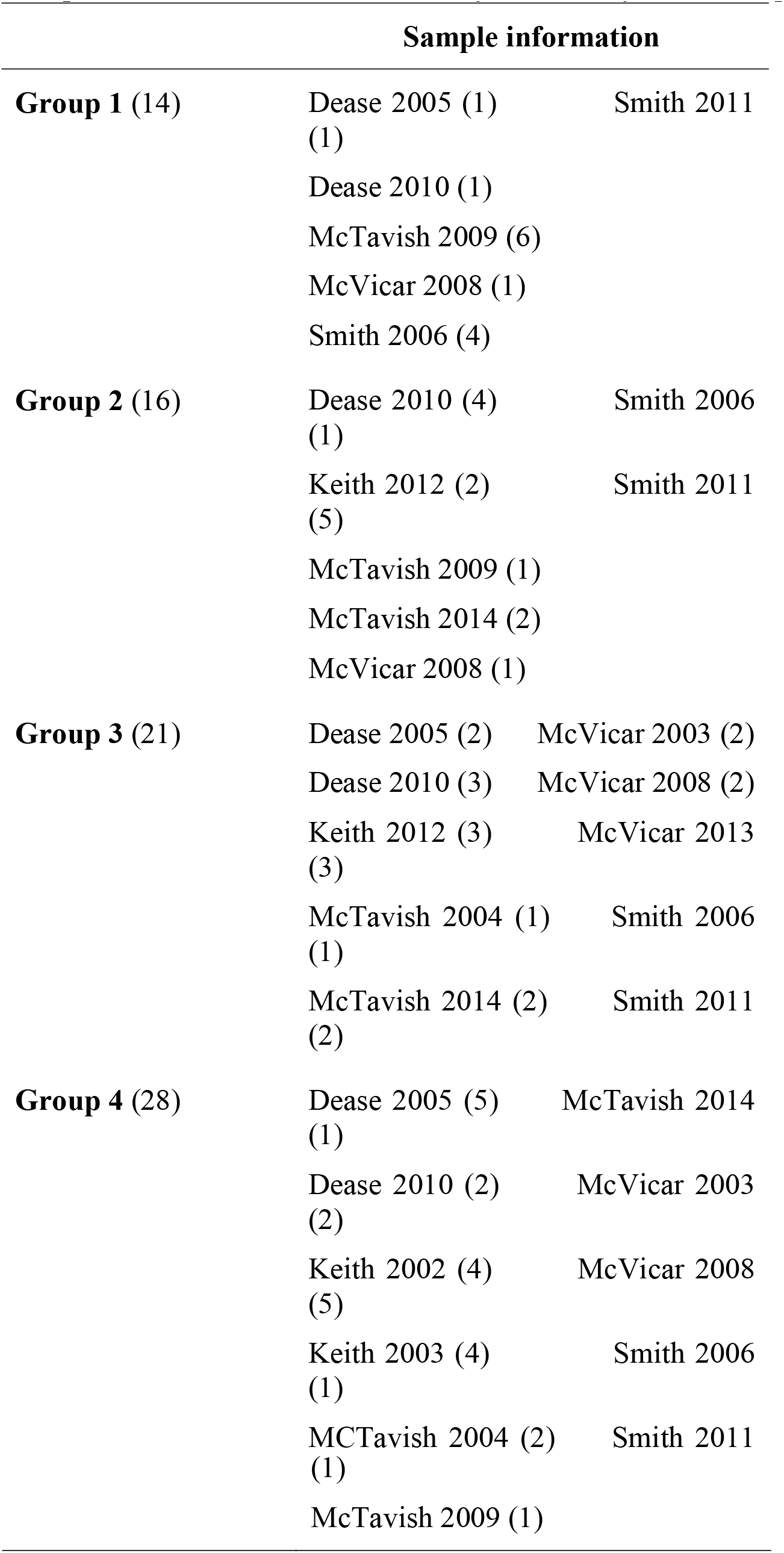
Spatial and temporal information for the 79 Lake Trout classified as piscivorous morph from Great Bear Lake and analyzed for fatty acids. Sample sizes are in brackets.

**Table A2.** Bayesian clustering (i.e., STRUCTURE, Pritchard et al. 2000) results for piscivorous morphotypes of lake trout from Great Bear Lake assessed using variation at 19 microsatellite markers. Shown are the mean log likelihood values (LnP[D]) for different hypothesized numbers of genetic populations (K) and the mean value of ΔK statistic of Evanno et al. (2005). Bold values represent the most likely number of genetic groups indicated by ΔK. Dashes = not applicable given that ΔK cannot be calculated for these values of K. For all STRUCTURE analyses, we employed an admixture model with the LOCPRIOR algorithm, correlated allelic frequencies, 100,000 burn-in and MCMC iterations and 10 iterations per K value were completed.

| K | Reps | Mean<br>LnP(D) | Delta<br>K |
| --- | --- | --- | --- |
| 1 | 10 | -10271.83 | — |
| 2 | 10 | **-10266.25** | **9.26** |
| 3 | 10 | -10572.68 | 0.03 |
| 4 | 10 | -10868.33 | 1.39 |
| 5 | 10 | -10739.53 | 0.45 |
| 6 | 10 | -10806.97 | 0.84 |
| 7 | 10 | -10678.37 | 0.66 |
| 8 | 10 | -10739.90 | 0.13 |
| 9 | 10 | -10862.98 | 1.09 |
| 10 | 10 | -10553.04 | — |

**Table A3.** Results of the discriminant analysis of principal components (DAPC, Jombart et al. 2010) implemented in the Adegenet package (Jombart et al. 2008) to determine the most likely number of genetic clusters (K) within the piscivorous Lake Trout form Great Bear Lake. The number of groups was identified using the find.clusters function (a sequential K-means clustering algorithm) and subsequent Bayesian Information Criterion (BIC), as suggested by Jombart et al. (2010). Stratified cross-validation carried out with the function *xvalDapc* was employed to determine the optimal number of PCs to retain in the analysis.

| **K** | **BIC** |
| --- | --- |
| 1 | 185.98 |
| 2 | 185.42 |
| 3 | 185.89 |
| 4 | 186.51 |
| 5 | 187.40 |
| 6 | 189.10 |
| 7 | 190.64 |
| 8 | 191.99 |
| 9 | 193.61 |
| 10 | 195.67 |

**Table A4.** Microsatellite loci used in this study and F_is_ values for each group per locus.

| **Locus** | **Group 1** | **Group 2** | **Group 3** | **Group 4** | **Giant** |
| --- | --- | --- | --- | --- | --- |
| **OtsG83b** | -0.021 | 0.017 | 0.046 | 0.090 | -0.013 |
| **Sco215** | 0.061 | 0.042 | -0.011 | -0.029 | 0.071 |
| **Smm17** | -0.433 | -0.069 | -0.038 | -0.304 | -0.012 |
| **Smm21** | -0.143 | -0.286 | -0.266 | 0.023 | 0.028 |
| **SnaMSU1** | 0.012 | -0.069 | -0.024 | -0.028 | -0.074 |
| **SnaMSU8** | -0.031 | 0.002 | 0.048 | 0.023 | 0.081 |
| **OMM1105** | 0.094 | -0.075 | -0.041 | -0.098 | -0.065 |
| **Smm22** | -0.014 | 0.055 | 0.082 | -0.088 | 0.137 |
| **SnaMSU13** | -0.105 | 0.053 | 0.136 | -0.073 | -0.049 |
| **SnaMSU5** | 0.088 | 0.065 | -0.032 | 0.039 | 0.159 |
| **Sco19** | -0.082 | 0.190 | 0.067 | -0.009 | 0.051 |
| **Sco202** | 0.107 | -0.166 | 0.047 | 0.115 | -0.080 |
| **SnaMSU10** | -0.108 | -0.030 | 0.086 | 0.096 | 0.203 |
| **SnaMSU12** | 0.122 | 0.069 | 0.123 | 0.201 | 0.072 |
| **SnaMSU6** | 0.008 | -0.090 | 0.002 | 0.007 | 0.207 |
| **Sal38** | -0.056 | 0.121 | -0.016 | -0.012 | 0.041 |
| **Sco200** | -0.015 | 0.098 | -0.096 | 0.041 | 0.244 |
| **SnaMSU11** | -0.060 | -0.108 | -0.012 | 0.083 | -0.011 |
| **SnaMSU3** | 0.059 | -0.019 | 0.065 | 0.012 | 0.085 |
| **Overall** | -0.027 | -0.011 | 0.009 | 0.005 | 0.057 |

**Fig. A1.**
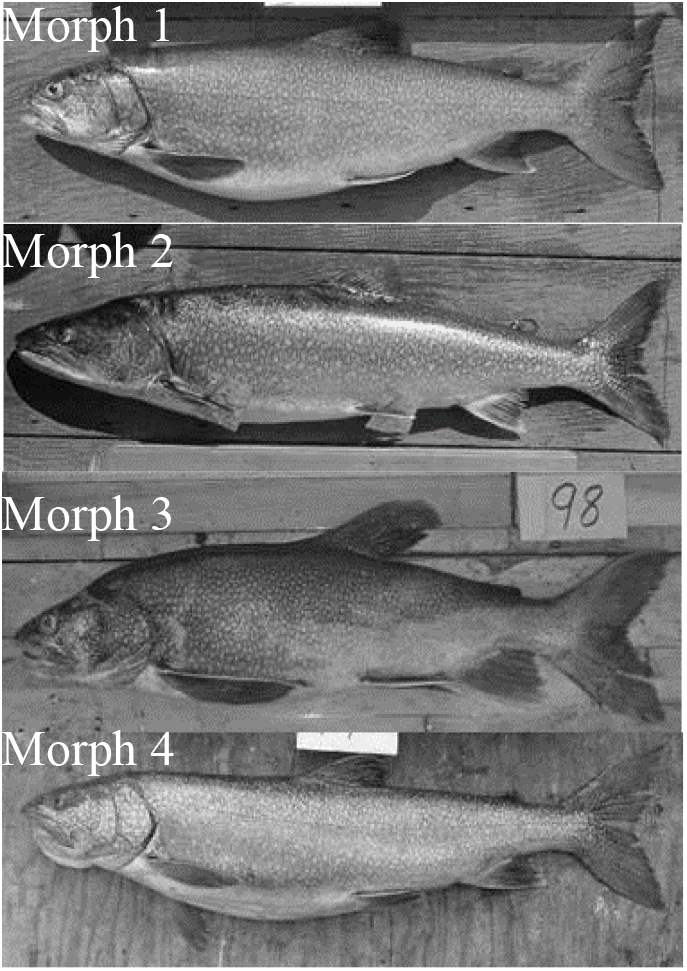
The four shallow-water morphotypes of Lake Trout from Great Bear Lake identified in Chavarie et al. (2013, 2015, 2016a, 2016b): the generalist, the piscivore, the benthic-oriented, and the pelagic specialist, Morphs1-4, respectively.

**Fig. A2.**
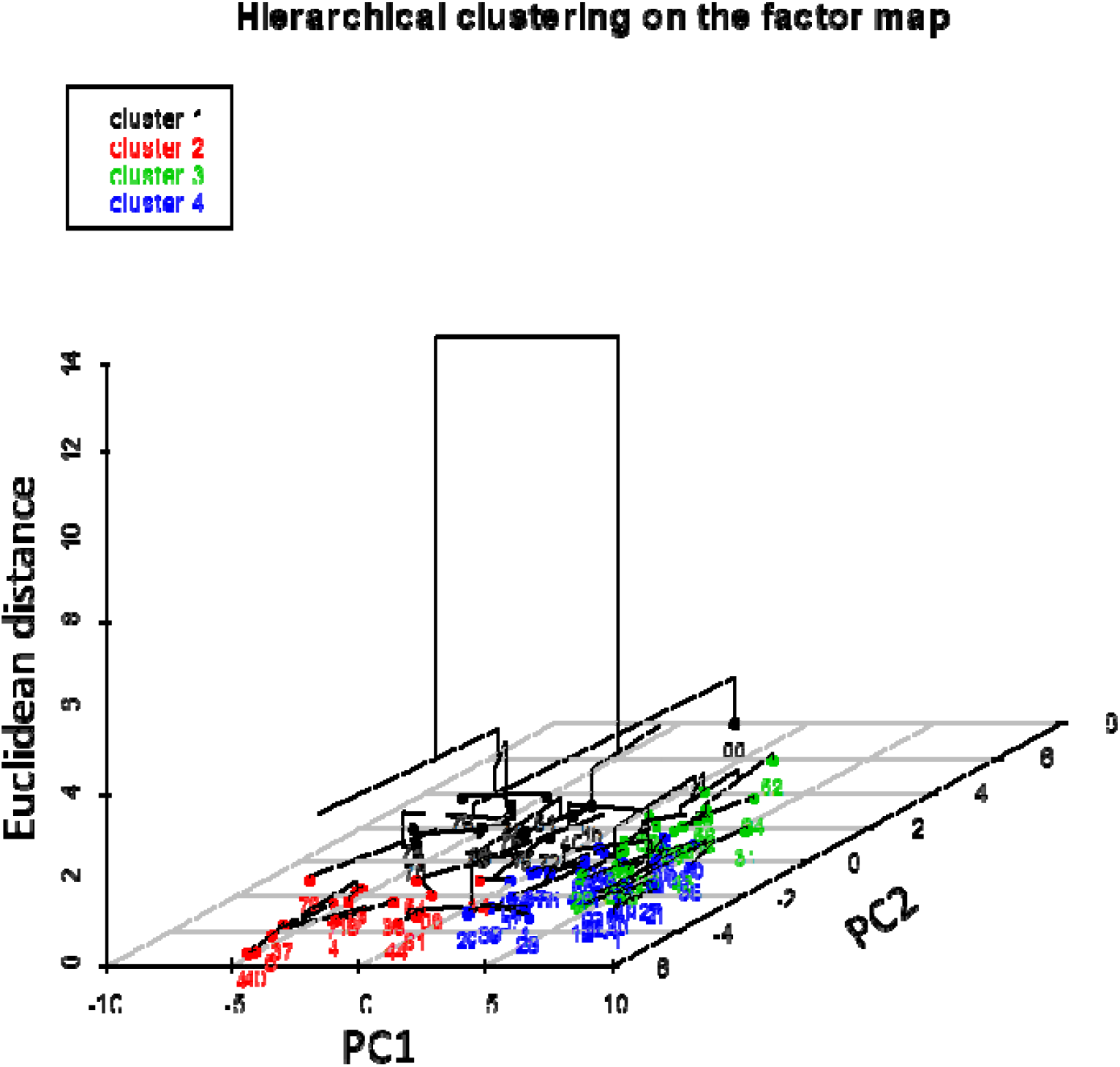
Hierarchical clusters of Great Bear Lake Lake Trout fatty acids profiles overlaid on the first two principal component axes (PCA) using FactoMineR.

**Fig. A3.**
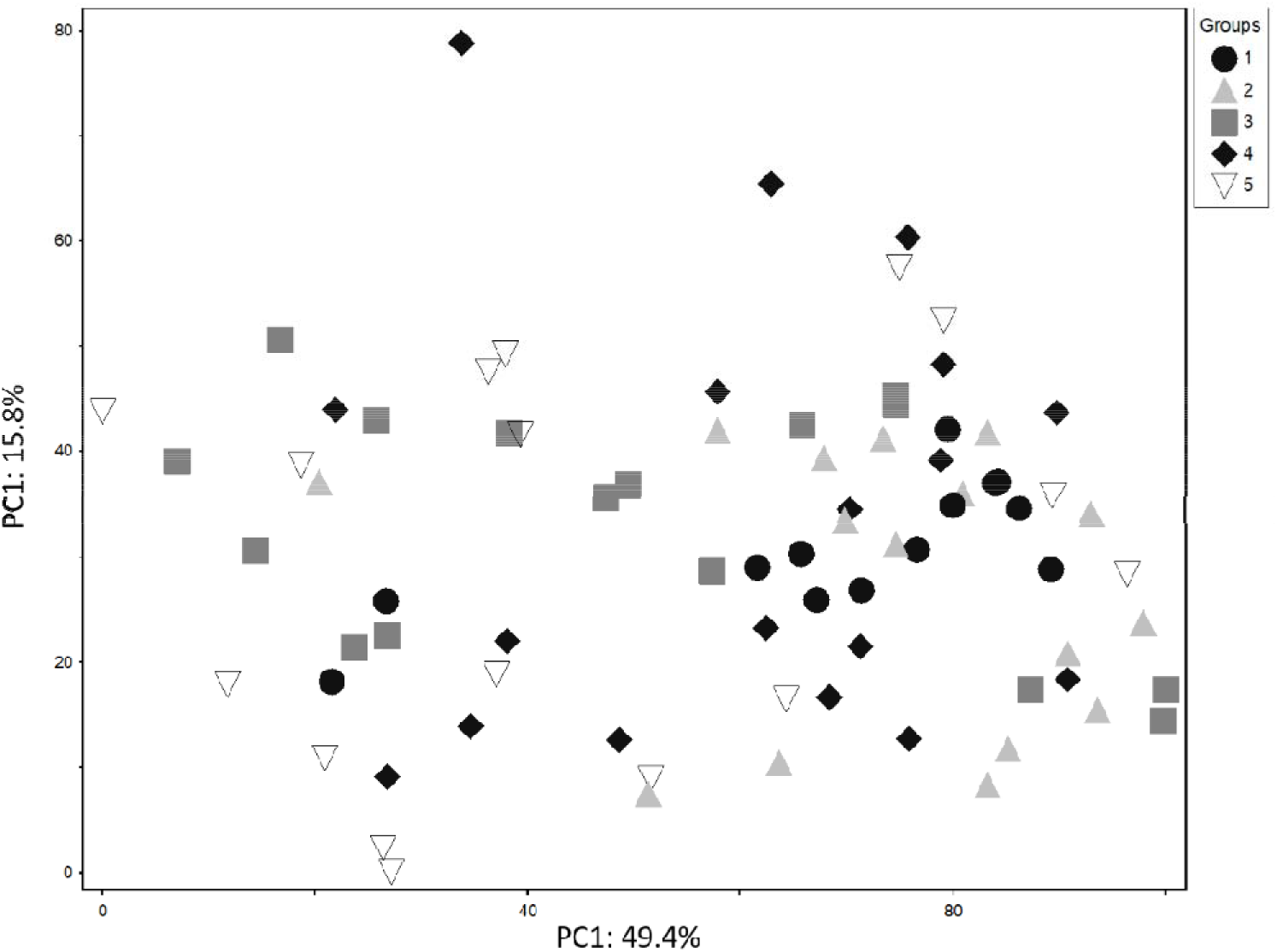
Principal Component Analysis (PCA) of fatty acids of 79 Lake Trout classified as piscivorous morph from Great Bear Lake, based on the proportions of 41 fatty acids in dorsal muscle tissue. Spatial variations (5 arms; 1=Keith, 2=McVicar, 3=McTavish, 4=Dease, and 5=Smith) are represented, based on the fatty acids profile of each lake trout analyzed in this study.

**Fig. A4.**
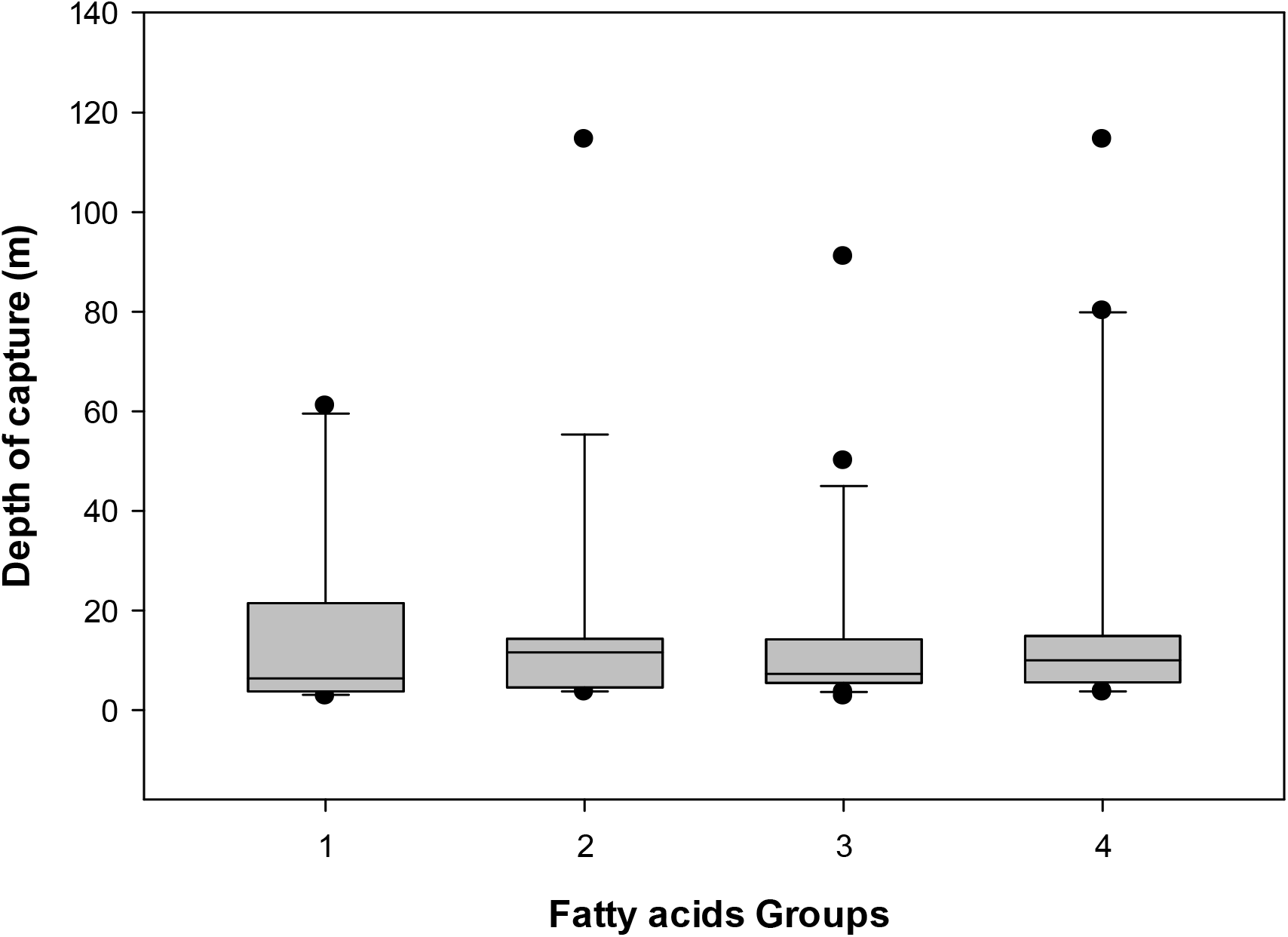
Depth of capture for four groups of piscivorous Lake Trout from Great Bear Lake (Groups identified by fatty acids profiles of individuals). Outliers are represented by a circle.

**Fig. A5.**
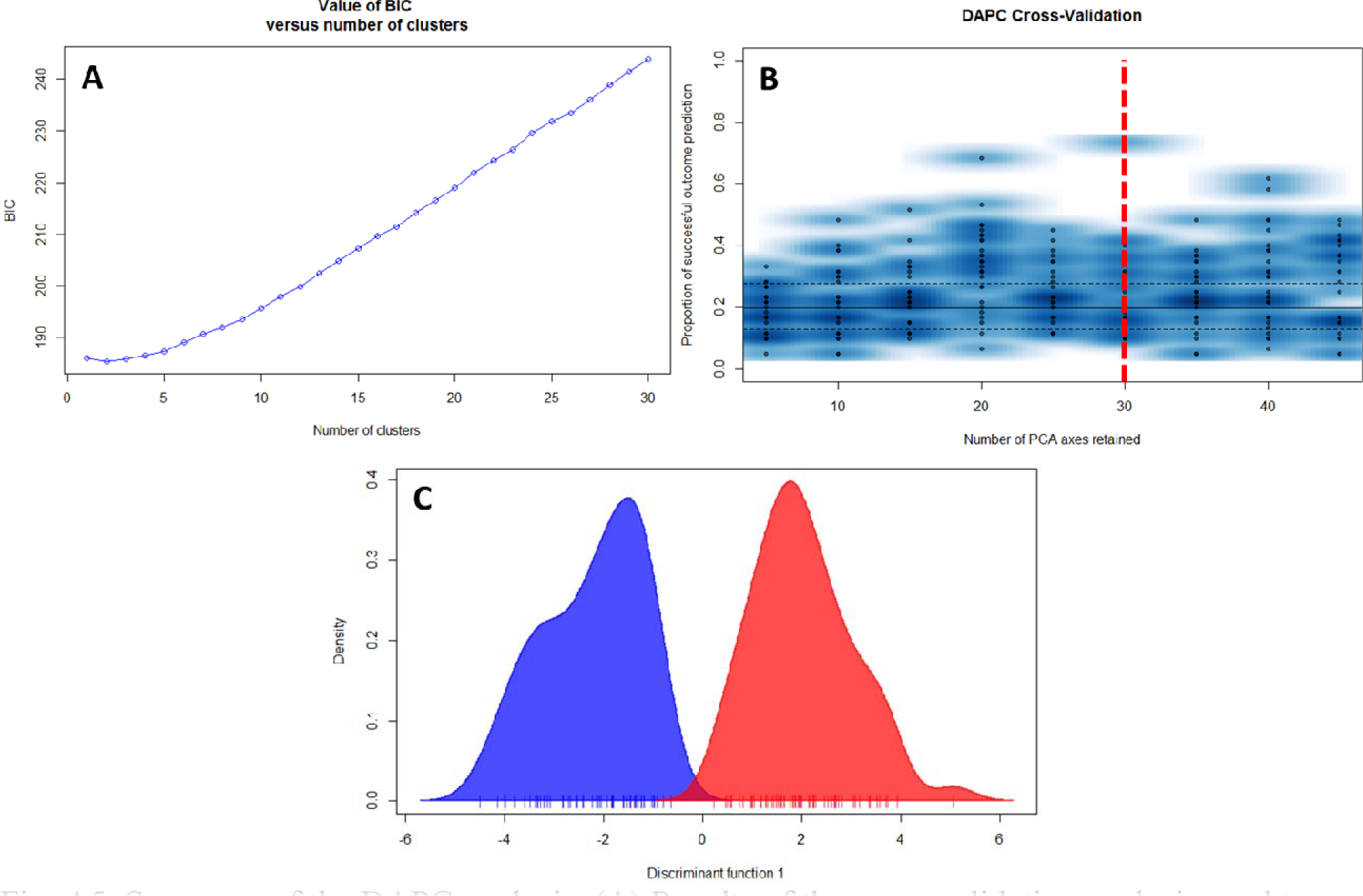
Summary of the DAPC analysis. (A) Results of the cross-validation analysis used to determine the number of PCs to retain in the DAPC analysis. Cross-validation analysis determined the most appropriate number of PCs retained was 30. (B) Inference of the number of clusters in the DAPC performed on piscivorous Lake Trout from Great Bear Lake. The function find.clusters was run with a maximum number of clusters of 10 to identify the optimal number of clusters based on the BIC values. A K value of 2 (the lowest BIC value) represents the best summary of the data (most probable number of (K)). (C) The results of the discriminant function that shows that the two clusters are mostly non-overlapping.

